# Elevated neuronal TAF15 expression leads to FTD-like phenotype

**DOI:** 10.64898/2025.12.13.694105

**Authors:** Tuo Yi, Haoyuan Guan, Jun Li, Wei Li, Bei Li, Caihong Zhu

## Abstract

TAF15, a DNA/RNA-binding protein whose dysfunction in alternative splicing impairs NMDA receptor signaling, can form amyloid fibrils that contribute to neurodegeneration in the neurodegenerative diseases like FTD and ALS, indicating a close involvement of TAF15 in both neuronal physiological function and activation of pathological pathways. However, the relationship between TAF15 expression levels and neurodegeneration, as well as the specific downstream pathways mediating its neurotoxicity, remains unclear. Here, we found a consistent upregulation of TAF15 in prefrontal cortex neurons from patients across multiple FTD and ALS subtypes. Both in vitro and in vivo experiments demonstrated that neuronal TAF15 overexpression triggered oxidative stress, evidenced by reactive oxygen species (ROS) production, leading to neurotoxicity and gliosis. Mice overexpressing TAF15 in medial prefrontal cortex (mPFC) neurons exhibited heightened anxiety and impaired fear memory, mimicking some key features of FTD behavioral abnormalities. Notably, these pathological and behavioral phenotypes are rescued by the antioxidant N-acetylcysteine amide (NACA), indicating that elevated TAF15 level exacerbates neurodegeneration primarily by activating oxidative stress pathways. Together, this work elucidates a novel TAF15-oxidative stress axis in FTD-associated neurodegeneration, providing a conceptual framework for future therapeutic development.

**Graphical abstract:** 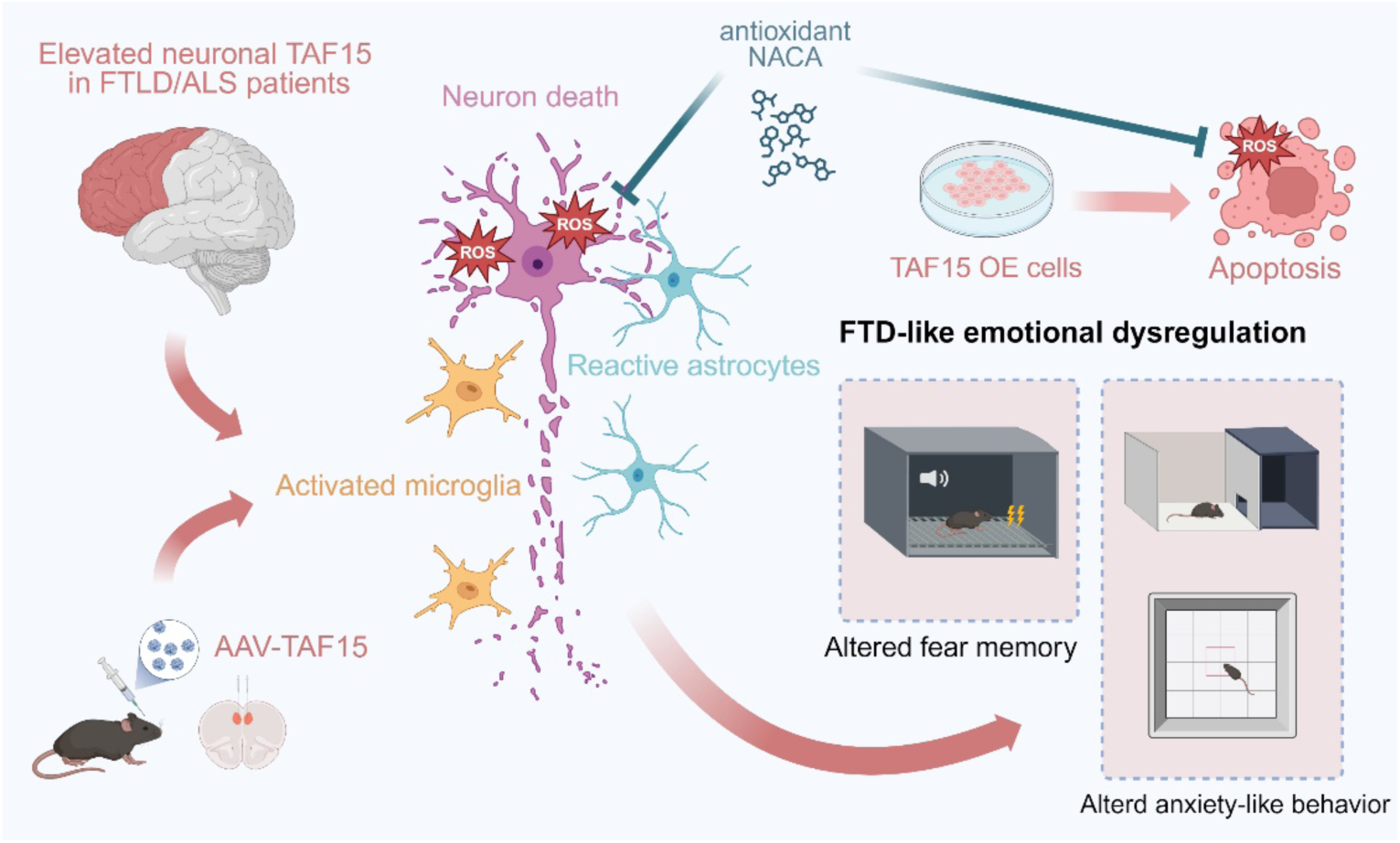

**Highlights:** TAF15 is upregulated in prefrontal cortex neurons of patients with FTD and ALS across different subtypes.

Elevated TAF15 expression provokes ROS elevation, leading to neurotoxicity and gliosis.

Neuronal TAF15 overexpression in the mouse mPFC results in FTD-like anxiety and impaired fear-memory phenotype.

Reducing oxidative stress can rescue FTD-like phenotype caused by TAF15 upregulation.

## Introduction

Frontotemporal dementia (FTD) is the second most common early-onset (<65 years) dementia worldwide, following Alzheimer’s disease^1^. Its high prevalence and the current lack of effective treatments impose a substantial societal burden. The symptoms of FTD are characterized by a range of personality changes and behavioral disturbances, including anxiety, apathy and other mood disorders, as well as a progressive decline in behavior and language^2–4^. Research on frontotemporal lobar degeneration (FTLD) has primarily focused on proteins such as TDP-43^5,6^, Tau^7–9^, and TMEM106B^10^. Recent studies, however, have revealed that approximately 10% of FTLD patients exhibit amyloid pathology characterized by aggregates of the FET (FUS, EWS, TAF15) protein family member TAF15 (TATA-box binding protein–associated factor 15) alone^11–14^. In these cases, TAF15 amyloid fibrils are predominantly localized within the cytoplasm of neurons, with minor presence in the neuron nucleus and glial cytoplasm^11^. Of note, TAF15 amyloid fibrils are not only implicated in FTLD but are also found within ascending and descending motor circuits, suggesting a potential role in the motor dysfunction phenotypes associated with diseases like amyotrophic lateral sclerosis (ALS)^15^. Indeed, missense mutations in *TAF15* have been reported in a subset of ALS patients; these variants form amyloid fibrils in vitro and induce neurodegeneration in Drosophila models^16–18^. For instance, the ALS-associated A31T mutation resides within the fibril core of TAF15 and forms stabilizing hydrogen bonds with the adjacent Q48 residue, thereby favoring the fibrillar conformation in vitro^19^. By contrast, to date no pathogenic *TAF15* coding mutations have been reported in FTD, where amyloid fibrils typically comprise wild-type TAF15^11^. Moreover, driven by its conserved N-terminal motifs, TAF15 can form homo- or hetero-aggregates with other FET family proteins; these assemblies are consistent with those observed in pathological models of neurodegenerative diseases such as ALS and FTD^20,21^.

TAF15, a DNA/RNA-binding protein, is predominantly nuclear but shuttles between the nucleus and cytoplasm to regulate transcription and RNA processing^22–25^. The N-terminal region (residues 1–201) of TAF15 contains a prion-like domain (PrLD), within which the SGYS motif is critical for driving the formation of amyloid fibrils^26^. Recent work further identified residues 7–99 within the N-terminal low-complexity domain (LCD) as a critical determinant of the fibrils isolated from FTD patient brains^11^. Beyond its role in amyloid pathology, TAF15, as an RNA-binding protein (RBP), is essential for the alternative splicing of RNAs involved in synaptic activity. For example, TAF15-mediated splicing is required for the synthesis of the zeta-1 subunit of the neuroplasticity-related Grin1 (NR1) receptor, thereby modulating the activity of NMDA glutamate receptors—dysregulation of which has been implicated in multiple neurodegenerative diseases, including ALS^27^. As a transcription factor, TAF15 overexpression has been shown to repress the transcription of *Npas4*, leading to reduced dendritic spine density in mouse hippocampal neurons and subsequent impairment of spatial memory^28,29^. Together, these findings underscore the importance of normal TAF15 function in maintaining neuronal physiology.

However, to date our understanding of TAF15 is still limited. The mechanisms by which TAF15 contributes to the pathogenesis of neurodegenerative diseases such as FTD and ALS, along with the specific downstream pathways it may elicit under pathological conditions, remains elusive. Here, we reported a significant correlation between elevated neuronal TAF15 expression and FTD and ALS. Using in vitro and in vivo TAF15 overexpression models, we revealed that elevated neuronal TAF15 activates oxidative stress, induces neurotoxicity and reactive gliosis. These pathological changes resulted in FTD-like behavioral phenotype in mice, including heightened anxiety and impaired fear memory. Notably, scavenging reactive oxygen species (ROS) with antioxidant N-acetylcysteine amide (NACA) rescued both the pathological and behavioral phenotypes caused by TAF15 overexpression, suggesting that oxidative stress represents a novel therapeutical target for TAF15-associated neurodegeneration. Overall, our study established a novel and valuable framework showing elevated TAF15 expression leads to FTD-like phenotype via promoting ROS production.

## Results

### Elevated neuronal TAF15 expression is associated with neurodegeneration and cytotoxicity

To test whether TAF15 expression levels are associated with the pathogenesis of neurodegenerative diseases, we analyzed snRNA-seq data from the prefrontal cortex of patients with FTLD and ALS across different subtypes (sporadic and C9orf72-linked). We found that TAF15 was significantly upregulated in neurons across these distinct disease subtypes, most prominently in inhibitory neurons (Figure 1A). This reveals a close relationship between elevated TAF15 expression and FTD/ALS spectrum. Thus, we sought to further elucidate the role of TAF15 upregulation in neurodegeneration by generating stable TAF15-overexpressing cell lines. Given that TAF15 contains an N-terminal low-complexity domain (LCD) and a C-terminal nuclear localization signal (NLS), and to preclude potential confounding effects from exogenously inserted FLAG tags, we constructed two full-length TAF15 overexpression models: one with a C-terminal FLAG tag (TAF15-FLAG OE) and another with an N-terminal FLAG tag (FLAG-TAF15 OE). Furthermore, since the N-terminal residues 7-99 of TAF15 have been reported to form amyloid fibrils in FTD patient brains, we also generated a cell line overexpressing this N-terminal fragment (TAF15 N-Term OE). Successful overexpression was confirmed at both the mRNA (Figure 1B, Figure S1A) and protein levels (Figure 1C-E; uncropped blots in Figure S2).

**Figure 1.**
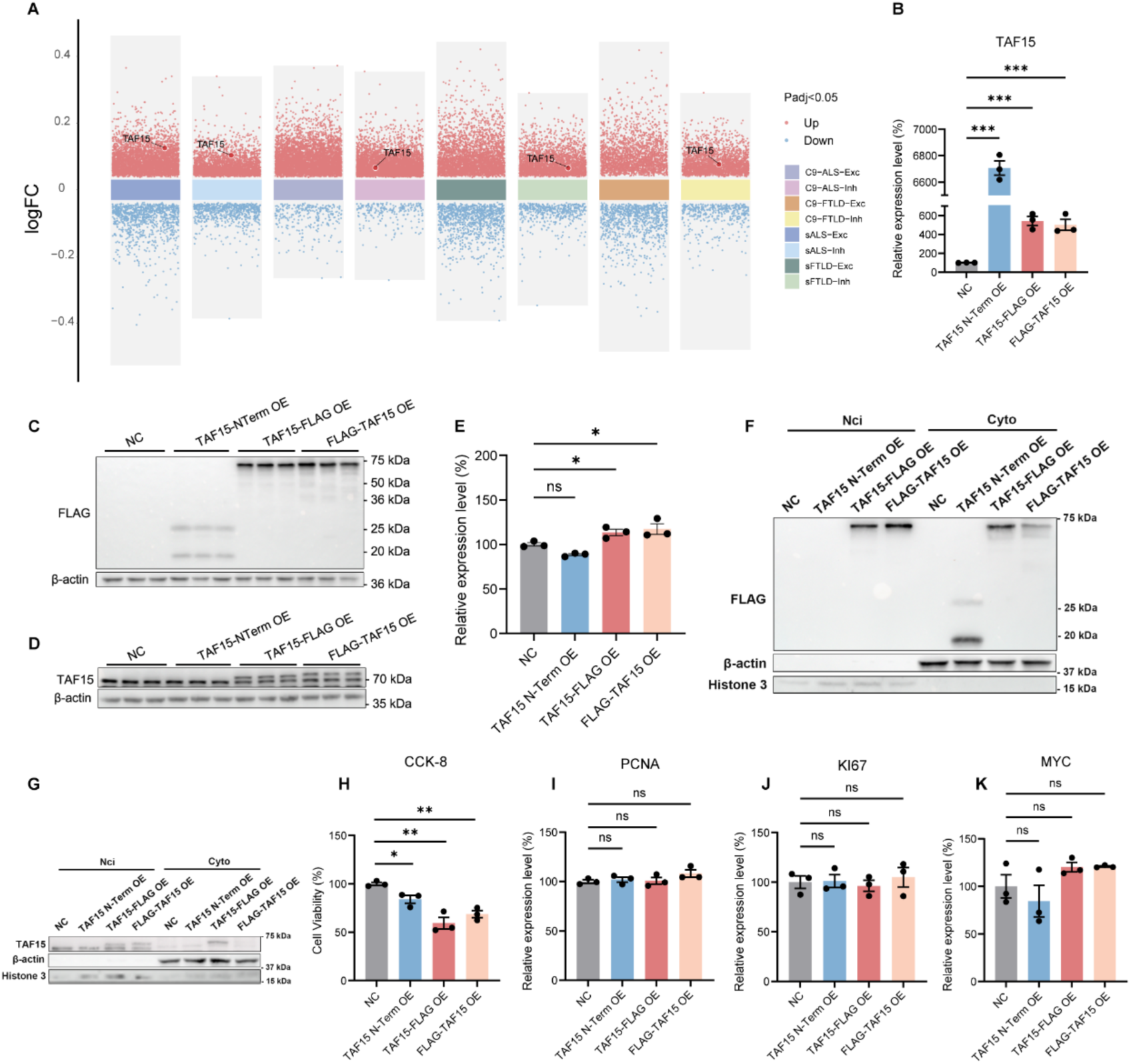
Elevated neuronal TAF15 expression is associated with neurodegeneration and cytotoxicity. (A) Multi-volcano plot of differentially expressed genes (DEGs) in frontal-cortex neuronal samples from ALS and FTLD patients across different subtypes. Upregulated genes are shown in red and downregulated genes in blue. The horizontal dashed line indicates the statistical significance threshold (adjusted *P* < 0.05). C9, C9orf72-linked; s, sporadic; Exc, excitatory neurons; Inh, inhibitory neurons. (B) Relative TAF15 mRNA expression in control mCherry-overexpressing cells, TAF15 N-term OE, TAF15-FLAG OE and FLAG-TAF15 OE cells (*n* = 3/group). Statistical comparisons were performed by one-way ANOVA with Dunnett’s post hoc test. (C) Immunoblot analysis of FLAG and β-actin (loading control) in the indicated cell lines (*n* = 3/group). (D) Immunoblot analysis of TAF15 and β-actin in the indicated cell lines (*n* = 3/group). (E) Quantification of TAF15 protein levels from (D). Statistical comparisons were performed by one-way ANOVA with Fisher’s LSD post hoc test. (F) Representative immunoblot of nuclear-cytoplasmic fractionation (loading ratio, nuclear : cytoplasmic = 1:15) probed for FLAG, β-actin (cytoplasmic marker), and Histone H3 (nuclear marker) in the indicated cell lines. (G) Representative immunoblot of nuclear-cytoplasmic fractionation (loading ratio, nuclear : cytoplasmic = 1:15) probed for TAF15, β-actin, and Histone H3 in the indicated cell lines. (H) Relative cell viability measured by CCK-8 assay in the indicated cell lines (*n* = 3/group). Statistical comparisons were performed by one-way ANOVA with Holm-Šidák post hoc test. (I-K) Relative mRNA expression levels of indicated proliferation-related genes in the indicated cell lines (*n* = 3/group). Statistical comparisons were performed by one-way ANOVA with Dunnett’s post hoc test.

Given prior reports that pathological TAF15 amyloid deposits in FTD localize to the neuronal cytoplasm rather than the nucleus, we performed nuclear–cytoplasmic fractionation to test whether overexpression drives nuclear export. As expected, the overexpressed TAF15 N-term fragment, which lacks the NLS, was exclusively localized to the cytoplasm (Figure 1F; uncropped blots in Figure S2). Although placement of the FLAG tag at the C-terminus modestly affected nuclear import (Figure 1F, G, Figure S1B, C; uncropped blots in Figure S2, S3), overall full-length TAF15 overexpression did not produce gross mislocalization. In line with the absence of cytoplasmic TAF15 mislocalization, Thioflavin-T (ThT) staining revealed no positive signal for amyloid fibrils in any of the overexpression models (Figure S1F). Given that TDP-43 pathology, characterized by nuclear clearing and cytoplasmic aggregation of phosphorylated TDP-43, is another hallmark of FTD and ALS, we assessed whether TAF15 overexpression induces TDP-43 pathology. Nuclear–cytoplasmic fractionation showed no TAF15-dependent TDP-43 nuclear depletion or change in cytosolic phosphorylated TDP-43 levels (Figure S1D, E; uncropped blots in Figure S3).

Having excluded mislocalization and canonical aggregation as primary consequences of TAF15 overexpression, we next asked whether TAF15 overexpression confers cellular toxicity. Using a CCK-8 cell viability assay, we found that overexpression of TAF15, particularly the full-length overexpression, significantly reduced cell viability (Figure 1H). Since SH-SY5Y is a neuroblastoma cell line, we verified that this effect was due to cytotoxicity rather than suppressed proliferation by examining the expression of common proliferation-related genes and found no change (Figure 1I-K). Taken together, we found that elevated neuronal expression of TAF15 is strongly associated with neurodegenerative diseases such as FTD and ALS. Cellular models featuring TAF15 overexpression exhibit cytotoxicity in the absence of amyloid fibril formation or TDP-43 pathology, suggesting that TAF15 overexpression may induce cytotoxicity via alternative downstream pathways.

### TAF15 upregulation disrupts gene expression related to oxidative stress and neuron viability

To explore the downstream pathogenic mechanisms of TAF15 upregulation cytotoxicity, we performed bulk RNA-Seq analysis on TAF15-FLAG OE cells. We found that TAF15 upregulation primarily activated pathways related to protein folding, oxidative stress, and apoptosis, while simultaneously suppressing pathways associated with axonogenesis, synapse assembly, and protein degradation mechanisms such as autophagy and the ubiquitin-proteasome system (Figure 2A, B). A protein–protein interaction (PPI) network analysis further illustrated the close relationships among these significantly altered pathways (Figure 2C). Additionally, Gene set enrichment analysis (GSEA) indicated substantial alterations in RNA splicing and mitochondrial gene expression pathways upon TAF15 overexpression (Figure 2D, E), which is consistent with the known role of TAF15 as a RBP and suggests a dysfunction in its alternative splicing capacity.

**Figure 2.**
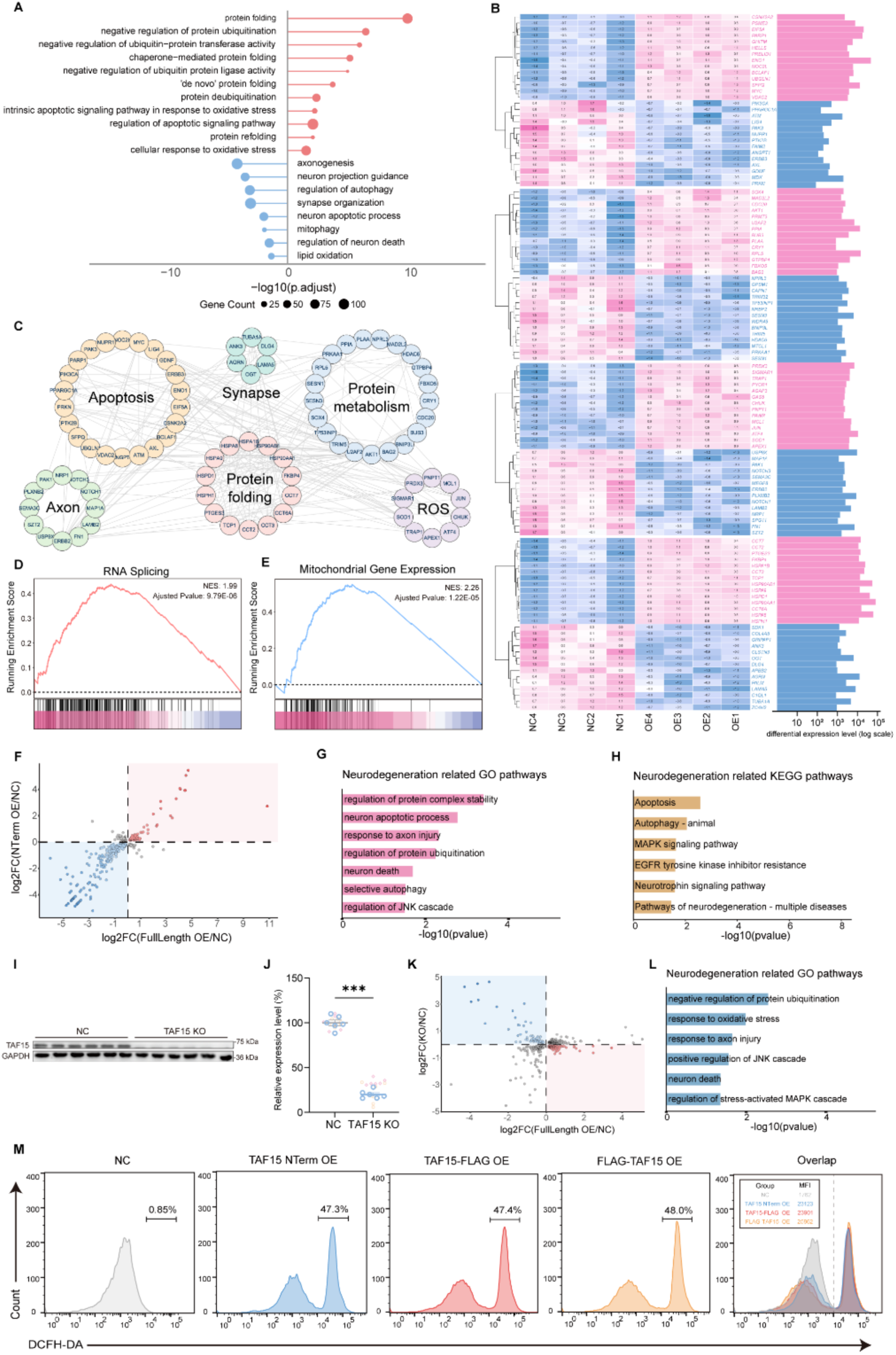
TAF15 upregulation disrupts gene expression related to oxidative stress and neuron viability. (A) GO enrichment analysis of significantly dysregulated pathways associated with neurodegeneration from DEGs in TAF15-FLAG OE cells. (B) Heatmap showing the top upregulated and downregulated genes implicated in neurodegeneration in TAF15-FLAG OE cells. (C) PPI network constructed from DEGs in TAF15-FLAG OE cells using STRING (interaction score ≥ 0.7) and visualized in Cytoscape. (D-E) GSEA plots of DEGs in in TAF15-FLAG OE cells. Adjusted *P* values and normalized enrichment scores (NES) are indicated. (F) Scatter plot comparing log₂FC of DEGs between in TAF15-FLAG OE cells (x-axis) and TAF15 N-term OE cells (y-axis). The region with concordant gene regulation is highlighted. (G) GO enrichment of commonly dysregulated pathways associated with neurodegeneration from the co-regulated genes shared between TAF15-FLAG OE and N-term OE cells. (H) KEGG pathway enrichment of the co-regulated genes shared between TAF15-FLAG OE and N-term OE cells, highlighting pathways relevant to neurodegeneration. (I) Immunoblot of TAF15 and GAPDH (loading control) in control and TAF15 knockout (KO) cells (*n* = 6/group). (J) Relative TAF15 mRNA expression in control and TAF15 KO cells (*n* = 6/group). Three independent replicate experiments were performed. (K) Scatter plot comparing log₂ fold-changes (log₂FC) of DEGs between TAF15 KO cells (x-axis) and TAF15 KO (y-axis) cells. The region showing opposite regulation between gain- and loss-of-function is highlighted. (L) GO enrichment analysis of commonly dysregulated pathways associated with neurodegeneration from the set of genes showing opposite expression trends in TAF15 KO cells versus KO cells. (M) Representative flow cytometry plots showing DCFH-DA fluorescence (indicating ROS levels) in the indicated cell lines after treatment with 10 μM DCFH-DA for 30 minutes.

To comprehensively delineate the key pathways downstream of TAF15 overexpression, we also conducted bulk RNA-Seq on TAF15 N-term OE cells. We found that the co-regulated genes in both overexpression models exhibited highly concordant expression trends (Figure 2F). Gene ontology (GO) enrichment analysis of these commonly regulated genes highlighted biological processes including protein complex homeostasis, neuronal apoptosis, and axon damage (Figure 2G). Corresponding KEGG pathway analysis further identified their involvement in apoptosis, autophagy, and MAPK signaling pathways (Figure 2H). Finally, to complement the gain-of-function data, we generated a TAF15 knockout cell line using CRISPR/Cas9 (Figure 2I, J; uncropped blots in Figure S3) and also performed bulk RNA-Seq. In comparison with the transcriptomic profiles, we identified a set of genes that exhibited inverse expression trends in the TAF15-FLAG OE and TAF15 KO cells (Figure 2K). GO enrichment analysis of these inversely regulated genes revealed significant enrichment in pathways such as oxidative stress response, axon damage and the negative regulation of ubiquitination (Figure 2L). To uncover the specific downstream mechanisms underlying TAF15-induced cytotoxicity, and, considering the above sequencing data and the well-documented activation of oxidative stress in nearly all neurodegenerative diseases, we measured ROS levels in the TAF15-overexpressing cells by fluorescent probe and flow cytometry. This revealed a significant increase in ROS levels across all three TAF15 overexpression models (Figure 2M), indicating that TAF15 upregulation induces cytotoxicity at least in part via the activation of oxidative stress pathways. Together, these transcriptomic datasets indicate that TAF15 dysregulation perturbs proteostasis, mitochondrial function, and oxidative stress responses - pathways plausibly linking TAF15 overexpression to neuronal dysfunction and cell death.

### TAF15 overexpression in mouse mPFC produces neurotoxicity and gliosis, resulting FTD-like phenotype

To investigate the pathogenic mechanisms of TAF15 upregulation in vivo and further understand its relevance to neurodegenerative diseases, we performed stereotactic injection of an AAV virus overexpressing TAF15 in neurons of the mouse medial prefrontal cortex (mPFC) (Figure 3A). Viral transduction produced robust TAF15 overexpression in the mPFC, and the overexpression signal co-localized with the AAV-expressed EGFP reporter (Figure 3B-C). Three weeks after injection, we subjected mice to a battery of behavioral assays to assess whether TAF15 overexpression leads to behavioral alterations (Figure 3D). In the open field test (OFT), TAF15-overexpressing (OE) mice spent significantly less time in the central zone (Figure 3E), showed a strong trend toward fewer entries into the center (Figure 3F), and traveled a significantly shorter distance in the center without changes in total locomotion (Figure 3G, H). Consistently, in the light dark transition test (LDT) TAF15 OE mice made fewer entries into the light compartment (Figure 3I) and exhibited a trend toward reduced time spent in the light compartment (Figure 3J). These results indicate that TAF15 OE mice display pronounced pathological anxiety-like behavior, while their basic motor function remains intact. Furthermore, in the cued fear conditioning test (CFC), TAF15 OE mice showed significantly reduced freezing time during the cue-on phase with unchanged freezing during the cue-off phase (Figure 3K), suggesting deficit in fear memory retrieval and regulation. By contrast, neither the Y-maze spontaneous alternation test nor the novel object recognition test (NOR) revealed overt deficits in working memory in TAF15 OE mice (Figure 3L, M). Collectively, these findings demonstrate that TAF15 overexpression in the mPFC produces FTD-like affective dysregulation, characterized by impaired modulation of anxiety and fear, without affecting general locomotion or working memory.

**Figure 3.**
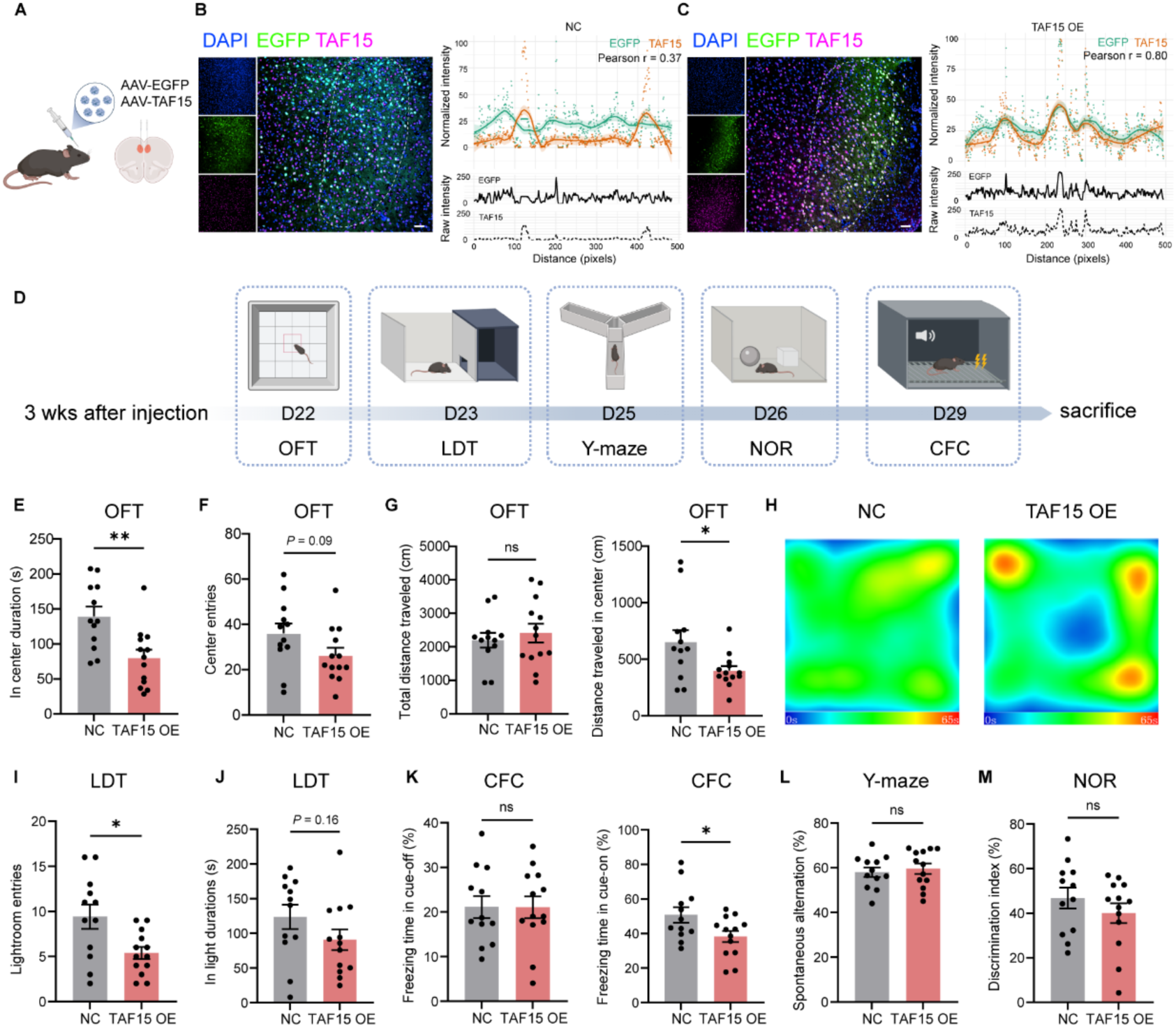
TAF15 upregulation in the mPFC causes FTD-like emotional dysregulation. (A) Schematic of AAV-mediated TAF15 overexpression in the mouse mPFC. (B) Colocalization profile of EGFP and TAF15 along a distance axis in mice stereotactically injected with control AAV. The normalized-intensity plot shows LOESS-smoothed fluorescence intensity for EGFP and TAF15 with the shaded area indicating the 95% confidence interval (CI). The raw intensity plot (y-axis fixed 0–250 for both channels) permits direct comparison of absolute signal across conditions. Data are presented as line plots with distinct line types for each channel. Scale bar, 50 μm. (C) Colocalization profile of EGFP and TAF15 along a distance axis in mice stereotactically injected with AAV-TAF15 (TAF15 OE). Scale bar, 50 μm. (D) Schematic timeline of behavioral tests. (E-G) mPFC-related anxiety behavior assessed by the open field test (*n* = 12 NC, *n* = 13 TAF15 OE). (H) Representative open-field trajectory heatmaps for control and TAF15 OE mice. (I-J) Anxiety behavior assessed by the light dark transition test (*n* = 12 NC, *n* = 13 TAF15 OE). (K) Fear memory regulation assessed by cued fear conditioning (*n* = 12 NC, *n* = 13 TAF15 OE). (L) Spatial working memory assessed by Y-maze spontaneous alternation test (*n* = 12 NC, *n* = 13 TAF15 OE). (M) Recognition working memory assessed by the novel object recognition test (*n* = 12 NC, *n* = 13 TAF15 OE).

To define neuropathological correlates of the behavioral phenotypes, we performed immunofluorescence analyses on the mPFC of TAF15 OE mice. We confirmed robust TAF15 upregulation in virus-transduced regions (Figure 4A). Consistent with in vitro experiments, we found that the expression of inducible nitric oxide synthase (iNOS), a marker for neuronal ROS production, was upregulated in TAF15 OE mice (Figure 4B), indicating in vivo activation of oxidative stress pathways. We next asked whether this oxidative stress resulted in neurotoxicity. Correspondingly, TUNEL staining revealed abundant apoptotic cells in TAF15 OE mice (Figure 4C), and NeuN staining showed a decreasing trend in neuronal density upon TAF15 overexpression (Figure 4D). Of note, TUNEL-positive signals were observed not only in neurons but also in surrounding glial-like cells (Figure 4C), prompting assessment of glial responses. In line with reactive gliosis observed in human neurodegenerative patients, TAF15 overexpression resulted in significant activation and proliferation of both microglia and astrocytes (Figure 4E, F). In line with in vitro experiments, TAF15 OE mouse brains revealed no detectable amyloid fibril formation by Congo red staining (figure S1G). Taken together, these in vivo results demonstrate that TAF15 upregulation induces oxidative stress, promotes excessive ROS production, and leads to aberrant apoptosis and gliosis in the brain.

**Figure 4.**
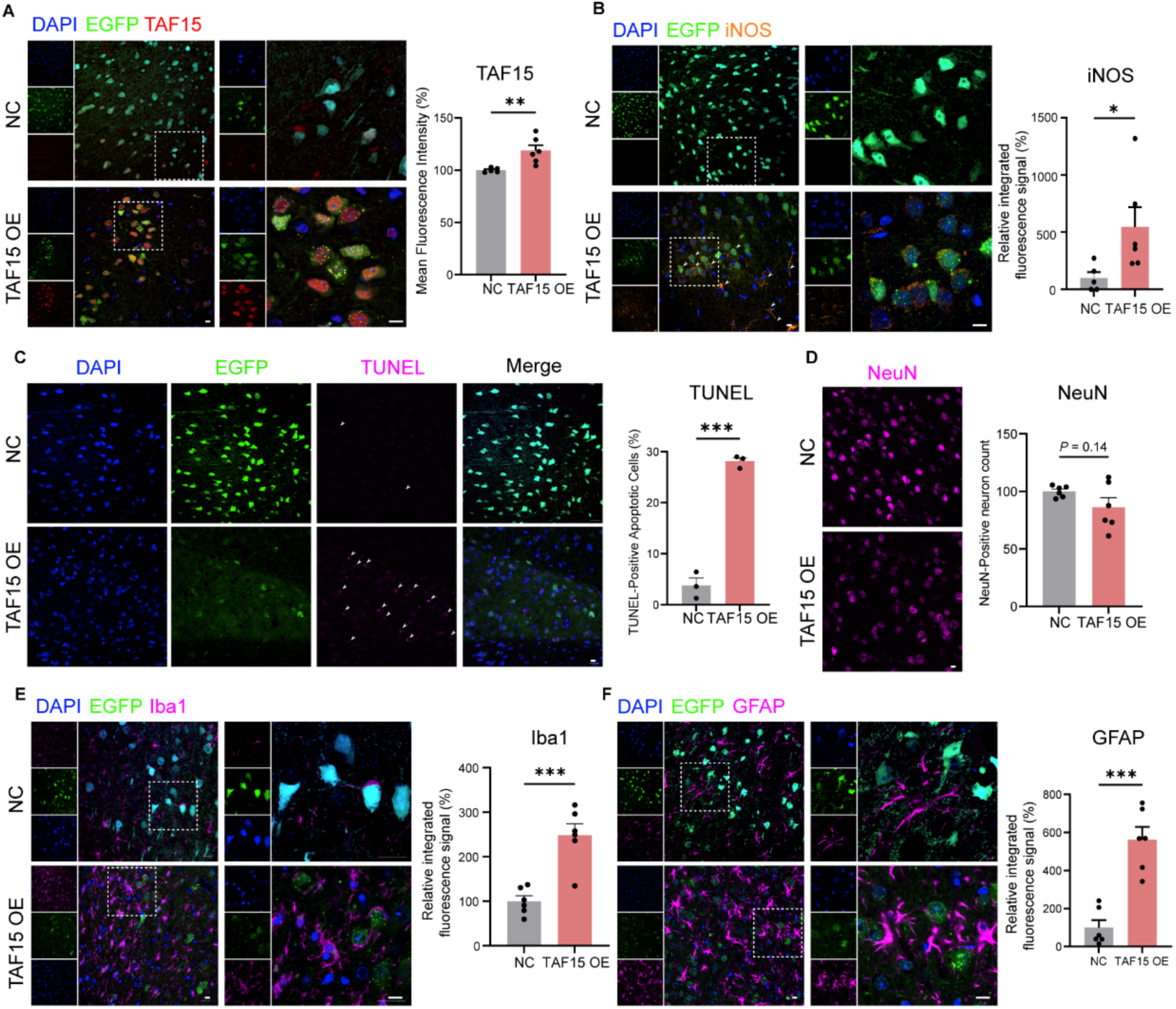
TAF15 upregulation in the mPFC induces ROS production, neurotoxicity and gliosis. (A) Mean fluorescence intensity (MFI) of TAF15 in the mPFC of NC and TAF15 OE mice (*n* = 6/group). Nuclei were counterstained with DAPI; EGFP marks viral transduction. Scale bar, 20 μm. (B) Integrated fluorescence signal of neuronal iNOS in the mPFC of NC and TAF15 OE mice (*n* = 6/group). Scale bar, 20 µm. (C) Quantification of apoptotic cells via TUNEL staining in the mPFC of NC and TAF15 OE mice (*n* = 3/group). Scale bar, 20 µm. (D) Quantification of neuronal density in the mPFC of NC and TAF15 OE mice (*n* = 6/group). Scale bar, 20 µm. (E) Quantification of microglial cell number in the mPFC of NC and TAF15 OE mice (*n* = 6/group). Scale bar, 20 µm. (F) Quantification of astrocyte number in the mPFC of NC and TAF15 OE mice (*n* = 6/group). Scale bar, 20 µm.

### Reducing oxidative stress can rescue FTD-like pathology caused by TAF15 upregulation

To test whether the FTD-like pathological phenotypes induced by TAF15 overexpression are mediated by oxidative stress, we first performed in vitro rescue experiments in TAF15-overexpressing cell lines using NACA, a blood-brain-barrier-permeable thiol antioxidant. NACA treatment reduced intracellular ROS levels in TAF15-overexpressing cells (Figure 5A), and notably, cell viability was largely restored to control levels upon NACA administration as measured by CCK-8 assay (Figure 5B).

**Figure 5.**
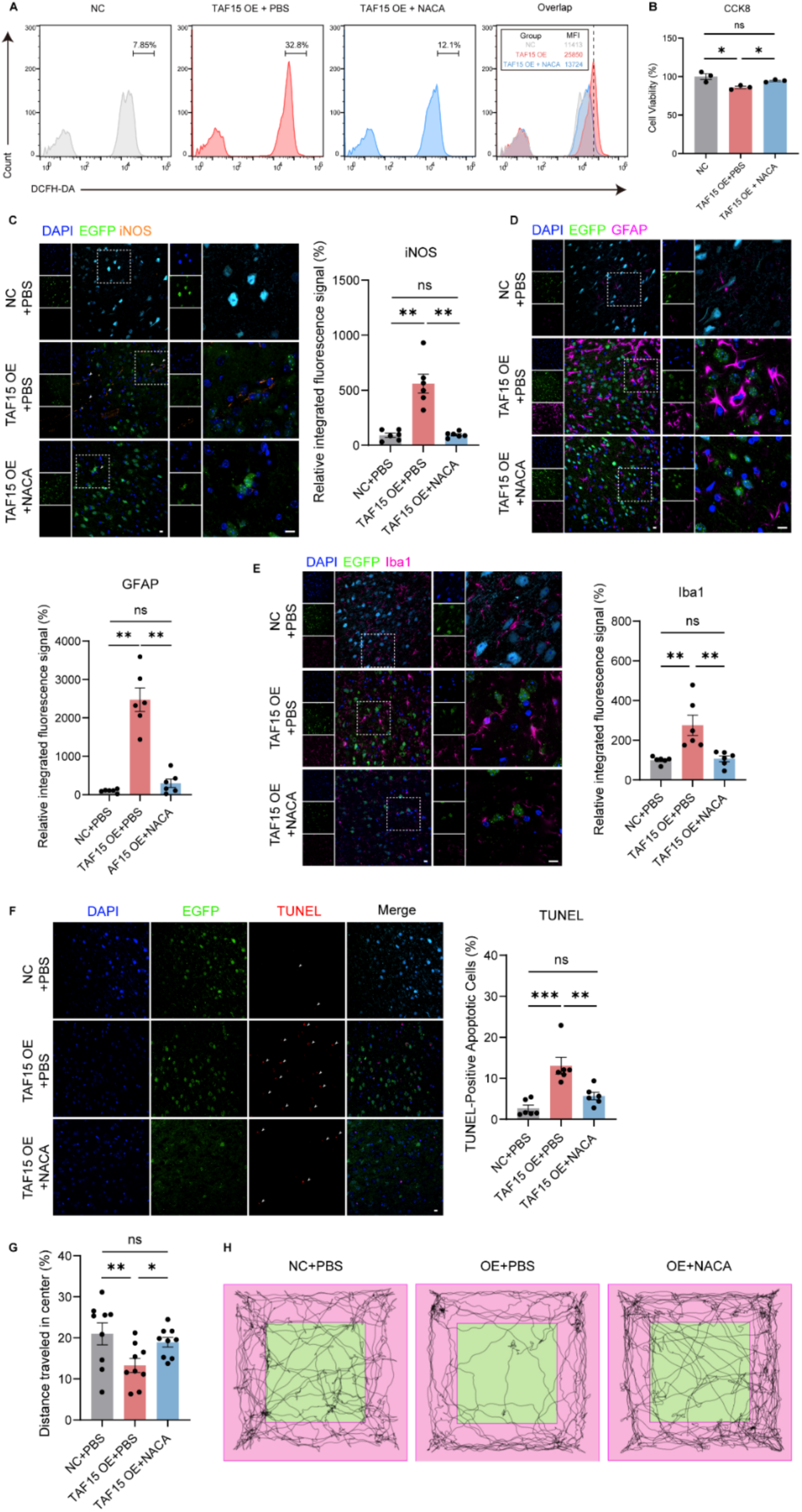
Reducing oxidative stress can rescue FTD-like pathology caused by TAF15 upregulation. (A) Representative flow cytometry plots of control, TAF15-FLAG OE, and TAF15-FLAG OE cells treated with NACA (250 μM, 24 hours), stained with 10 μM DCFH-DA for 30 minutes to measure ROS levels. (B) Relative cell viability measured by CCK-8 assay in control, TAF15-FLAG OE, and NACA-treated TAF15-FLAG OE cells (*n* = 3/group). Statistical comparison by one-way ANOVA with Fisher’s LSD post hoc test. (C) Integrated fluorescence signal of neuronal iNOS in the mPFC of control AAV-injected mice treated with PBS (i.p., *n* = 6), TAF15 OE mice treated with PBS (i.p., *n* = 6), and TAF15 OE mice treated with NACA (i.p., *n* = 6, 200 mg/kg). Viral transduction was indicated by EGFP. Scale bar, 20 μm. Statistical comparison by Brown-Forsythe and Welch ANOVA tests with Dunnett’s T3 post hoc test. (D) Quantification of astrocyte number in the mPFC of the indicated mice (*n* = 6/group). Scale bar, 20 μm. Statistical comparison by Brown-Forsythe and Welch ANOVA tests with Dunnett’s T3 post hoc test. (E) Quantification of microglial cell number in the mPFC of the indicated mice (*n* = 6/group). Scale bar, 20 μm. Statistical comparison by one-way ANOVA with Tukey’s HSD post hoc test. (F) Quantification of apoptotic cells via TUNEL staining in the mPFC of the indicated mice (*n* = 6/group). Scale bar, 20 μm. Statistical comparison by one-way ANOVA with Tukey’s HSD post hoc test. (G) Open field test assessing anxiety-like behavior in the indicated mice (*n* = 9/group). Statistical comparison by one-way ANOVA with Fisher’s LSD post hoc test. (H) Representative open-field trajectory heatmaps for the groups in (G).

We then evaluated the effects of systemic antioxidant treatment in vivo by administering NACA via intraperitoneal injection to TAF15 OE mice. Immunofluorescence analysis demonstrated that NACA significantly reduced iNOS expression levels, and thus attenuated ROS accumulation in the mPFC of TAF15 OE animals (Figure 5C). Concomitantly, the activation and proliferation of both astrocytes and microglia, which were prominent in TAF15 OE mice, were substantially ameliorated following NACA treatment (Figure 5D, E). TUNEL staining also revealed a significant reduction in apoptotic signals following NACA treatment, indicating that oxidative stress inhibition alleviates TAF15-induced cellular apoptosis (Figure 5F). Finally, to assess the pathological improvements translated into behavioral recovery, we subjected TAF15 OE mice to the open field test 24 hours after NACA treatment. We found that Intraperitoneal NACA significantly rescued the anxiety-like phenotype of TAF15 OE mice, returning measures of central zone exploration to levels comparable to control animals (Figure 5G, H). Together, these results reveal that pharmacological reduction of ROS by NACA attenuates neuronal apoptosis, gliosis and reverses FTD-like anxiety phenotypes, indicating that oxidative stress per se is a major mediator of TAF15-induced neurotoxicity in neurodegeneration diseases.

## Discussion

Neurodegenerative diseases such as Alzheimer’s disease (AD), FTD and ALS, are always associated with dysregulation of specific proteins, including the well-characterized Aβ^30^, Tau^7–9^, TDP-43^5,6^, and FUS^15,19^. Aberrant expression, hyperphosphorylation, or altered alternative splicing of these proteins often represent key pathogenic drivers in neurodegeneration^31–33^. With ongoing research, additional proteins implicated in neurodegenerative pathogenesis have been identified, including TAF15. However, the underlying mechanisms of these protein aggregates in neurodegeneration remain largely unknow. Importantly, a distinctive feature of FTLD cases with TAF15 pathology is its mutual exclusivity with the major pathological aggregates of TDP-43 (approximately 50% of FTLD cases) and Tau (approximately 40%), instead harboring amyloid fibrils formed by residues 7–99 of the N-terminal prion-like domain of TAF15^11,12^. This defines a pathological basis for the FTLD-FET subtype that comprises approximately 10% of FTLD cases^14^, suggesting that TAF15 may contribute - directly or indirectly - to the pathogenesis of FTD, ALS and related neurodegenerative disorders.

Research into the role of TAF15 in neurodegeneration remains at an early stage, with many underlying mechanisms yet to be elucidated. We first sought to determine whether TAF15 expression levels per se correlate with disease development. Analysis of snRNA-seq data from the prefrontal cortex of ALS and FTLD patients revealed robust upregulation of TAF15 in neurons across different disease subtypes (sporadic and C9orf72-linked), indicating a strong association between TAF15 expression and disease. As a canonical FET family member, TAF15 is primarily nuclear under physiological conditions^21^. However, pathological TAF15 amyloid fibrils are primarily observed in the neuronal cytoplasm. Whether cytoplasmic mislocalization precedes fibril formation or, conversely, fibrillization occurs before nuclear import remains unresolved. Our findings demonstrate that TAF15 overexpression alone does not cause its mislocalization from the nucleus nor directly induce amyloid fibril formation. Although mere overexpression is insufficient, previous reports indicate that under stress conditions such as heat shock, overexpressed TAF15 is recruited into stress granules^34^. Notably, chronic stress or proteostatic imbalance can facilitate the transition of these dynamic assemblies into persistent, pathological amyloid fibrils^35–37^. Integrated bulk RNA-seq analysis of cells overexpressing either full-length or N-terminal TAF15, as well as TAF15 knockout cells, revealed not only activation of pathways related to oxidative stress, apoptosis and axonal damage but also significant TAF15 involvement in pathways governing protein folding, particularly those involving heat shock proteins (HSPs). This aligns with reports of prominent HSP upregulation in the brains of patients with various neurodegenerative diseases, including AD, FTLD, and ALS^38,39^. Concurrently, we observed dysregulation of protein degradation pathways, such as autophagy-lysosome and ubiquitin-proteasome pathway, indicating that TAF15 overexpression broadly disrupts proteostasis, from folding to degradation.

To investigate in vivo consequences of TAF15 upregulation, we used AAV-mediated stereotaxic delivery to overexpress TAF15 in neurons of mPFC where the main pathological changes were observed in FTD patient. Patients and mouse models with FTD often exhibit various psychological disturbances and cognitive decline, including heightened anxiety, apathy, and impaired fear memory processing^40–44^. In line with that, mice with mPFC neuronal TAF15 overexpression displayed anxiety-like behaviors and dysregulated fear memory reminiscent of FTD-like phenotype in behavioral assays (OFT, LDT, CFC). Bioinformatic and pathological analyses revealed significant activation of oxidative stress in both in vitro and in vivo TAF15 overexpression models. Importantly, treated with the antioxidant NACA to scavenge ROS and mitigate oxidative stress substantially rescued the neurotoxicity, pathological changes, and behavioral phenotypes induced by TAF15 overexpression. Physiologically, moderate ROS levels are integral to cell signaling and mitochondrial dynamics^45^. However, impaired electron transport, altered membrane potential, or defective clearance mechanisms lead to sustained ROS elevation that damages macromolecules (DNA, lipids and proteins) and ultimately precipitates cell death^46^. The brain, with its high oxygen consumption and lipid-rich environment, is particularly vulnerable to oxidative stress, which constitutes a key pathological feature across neurodegenerative diseases^46^. Our study therefore suggests a close relationship between TAF15 upregulation and ROS activation, unveiling a novel TAF15-mediated pathogenic mechanism in neurodegeneration diseases. Furthermore, TAF15 overexpression in the mouse mPFC not only induced neuronal apoptosis but also prompted the activation and proliferation of astrocytes and microglia, consistent with observations reported in brains of patients with the aFTLD-U (FTLD-FET) subtype^47^. Glial cells perform essential roles in neuronal support and regulation, clearing waste and maintaining homeostasis^48–50^. However, chronic gliosis leads to the release of pro-inflammatory cytokines and chemokines, fostering a state of chronic neuroinflammation that exacerbates neuronal damage and apoptosis, thus accelerating neurodegeneration^48,51,52^. Therefore, activation and proliferation of glial cells may further exacerbate TAF15-mediated neurodegeneration.

## Limitations of the study

In this study, we established a framework showing that TAF15 overexpression leads to FTD-like phenotype by activating oxidative stress pathways. Nonetheless, some key mechanistic questions remain - most notably, how TAF15 drives ROS generation and how neuronal TAF15 upregulation elicits glial activation without inducing amyloid fibril formation or TDP-43 pathology. As a transcription factor and RNA-binding protein, TAF15 dysregulation may alter the expression of crucial genes/pathways involved in mitochondrial functions. In line with that, our GO enrichment and GSEA analyses indicated substantial alterations in mitochondrial function pathways upon TAF15 overexpression. Given that mitochondria are the primary source of intracellular ROS, mitochondrial dysfunction could lead to substantial ROS accumulation^53^. Neuron-to-glia signaling may also be mediated by damage-associated molecular patterns (DAMPs) released from TAF15-induced apoptotic neurons and recognized by glial pattern recognition receptors (e.g., Toll-like receptors), precipitating innate immune activation and thus driving glial activation and proliferation^54–56^. The absence of amyloid fibril formation may be attributed to the intrinsic limitations of models we used. Similarly, a recent study demonstrated that transgenically overexpressing of the lysosomal transmembrane 106B (TMEM106B) in mice leads to synaptic function and neuronal health without amyloid fibril formation^57^. Therefore, these studies suggest a dissociation between amyloid fibril formation and neurotoxicity/neurodegeneration. Alternatively, long-term overexpression of TAF15 in mice is warranted to determine whether amyloid fibril formation is time-dependent, although this is not the case in the TMEM106B study^57^. Together, future work dissecting these specific mechanisms and questions will be essential to evaluate TAF15 as a therapeutic target for related neurodegenerative disease.

## Resource availability

### Lead contact

Further information and requests should be directed to and will be fulfilled by the lead contact, Caihong Zhu.

## Materials availability

This study did not generate new reagents.

## Data and code availability

- Raw bulk RNA-seq data has been deposited in the NCBI Sequence Read Archive under BioProject accession PRJNA1376244. Raw snRNA-seq data used in this study are available from Synapse under accession syn51105515. Processed expression matrices and analysis scripts are available from the lead contact upon request.
- This paper does not report original code.
- Any additional information required to reanalyze the data reported in this paper is available from the lead contact upon request.

## Acknowledgments

C. Zhu is sponsored by Research Startup Funds of Fudan University, National Natural Science Foundation of China (No. 82271476 and No. 82071436), Shanghai Pujiang Program (No. 20PJ1401100), and The Program for Oriental Scholars of Shanghai Universities (Distinguished Professor)(No. TP2022050). B. Li is supported by National Natural Science Foundation of China (No. 82101502) and China Postdoctoral Science Foundation (No. 2021M690036). The funders had no role in study design, data collection and analysis, decision to publish, or preparation of the manuscript.

## Author contributions

Conceptualization, C.Z., B.L., T.Y.; methodology, C.Z., B.L., T.Y.; investigation, T.Y., H.G., J.L., W.L., Y.S., W.H.; writing—original draft, C.Z., B.L., T.Y.; writing—review and editing, C.Z., B.L., T.Y.; funding acquisition, C.Z., B.L.; supervision, C.Z., B.L.; resources, C.Z., B.L.

## Declaration of interests

The authors declare no competing interests.

## STAR★Methods

### Key resources table

| REAGENT OR RESOURCE | SOURCE | IDENTIFIER |
| --- | --- | --- |
| **Antibodies** |  |  |
| Rabbit monoclonal anti-NeuN | Abcam | ab177487 |
| Rabbit polyclonal anti-GFAP | Abcam | ab7260 |
| Rabbit monoclonal anti-Iba1 | FUJIFILM Wako | 019-19741 |
| Rabbit polyclonal anti-TAF15 N-terminal region | Aviva Systems Biology | ARP30112_P050 |
| Mouse monoclonal anti-TDP-43 | Proteintech | 60019-2-Ig |
| Rabbit monoclonal anti-phospho-TDP43 | Proteintech | 80007-1-RR |
| HRP-conjugated goat polyclonal anti-rabbit | Proteintech | SA00001-2 |
| HRP-conjugated goat polyclonal anti-mouse | Proteintech | SA00001-1 |
| Mouse monoclonal anti-β-actin | Proteintech | 66009-1-Ig |
| Mouse monoclonal anti-GAPDH | Proteintech | 60004-1-Ig |
| Mouse monoclonal anti-β-Tubulin | Abmart | M20005 |
| Rabbit monoclonal anti-iNOS | ABclonal | A3774 |
| Mouse monoclonal anti-DDDDK-Tag | ABclonal | AE005 |
| Rabbit monoclonal anti-Histone 3 | ABclonal | A17562 |
| Alexa Fluor™ 647-conjugated donkey polyclonal anti-rabbit | Thermofisher | A-31573 |
| **Plasmids and virus strains** |  |  |
| AAV9-EGFP | this study | N/A |
| AAV9-TAF15-3FLAG-EGFP | this study | N/A |
| lentiCRISPRv2 | this study | N/A |
| lentiCRISPRv2-TAF15 sgRNA | this study | N/A |
| pLV3-CMV-mCherry | this study | N/A |
| pLV3-CMV-TAF15-3×FLAG-mCherry | this study | N/A |
| pLV3-CMV-3×FLAG-TAF15-mCherry | this study | N/A |
| pLV3-CMV-TAF15 N-term-3×FLAG-mCherry | this study | N/A |
| **Critical chemicals and commercial assays** |  |  |
| NheI | NEB | R3131S |
| BamHI | NEB | R3136S |
| BsmBI-v2 | NEB | R0739S |
| DMEM | Gibco | 15596018 |
| DAPI | Sigma | D9542 |
| NACA | APExBIO | B8778 |
| Congo Red | APExBIO | B7766 |
| Carazzi's Hematoxylin Stain Solution | APExBIO | K2614 |
| Thioflavin T | absin | abs42237191 |
| DCFH-DA | MeilunBio | MB4682-1 |
| Fetal bovine serum | MeilunBio | PWL001-3 |
| Cell Counting Kit-8 | MeilunBio | MA0218 |
| BCA Protein Assay Kit | MeilunBio | MA0082-1 |
| TUNEL Apoptosis Assay Kit | YEASEN | 40308ES50 |
| HighGene transfection reagent | ABclonal | RM09014P |
| EZ-press RNA Purification Kit | EZBioScience | B0004D |
| 2×Color SYBR Green qPCR Master Mix(ROX2) | EZBioScience | A0012-R2 |
| 4×EZscript Reverse Transcription<br>Mix II (with gDNA Remover) | EZBioScience | EZB-RT2GQ |
| **Experimental models:**<br>**Organisms/strains** |  |  |
| Mouse: Wild type (C57BL/6J) | Cyagen | N/A |
| SH-SY5Y cell line | ATCC | CRL-2266 |
| 293T cell line | ATCC | CRL-3216 |
| **Software and algorithms** |  |  |
| R 4.5.2 | R Core Team | https://cran.r-project.org/ |
| ImageJ | NIH Image | https://imagej.en.softonic.com/ |
| FlowJo™ v10.8.1 | BD Biosciences (previously Tree Star) | https://flowjo.com/flowjo10/download |
| Graphpad Prism v10.1.2 | GraphPad | https://www.graphpad.com/ |
| Graphical design software | BioRender | https://biorender.com/ |

### Experimental model and study participant details Cells

Human neuroblastoma SH-SY5Y cells were cultured at 37°C in a humidified atmosphere with 5% CO₂ in DMEM supplemented with 10% (v/v) Fetal bovine serum (FBS) and 1% (v/v) penicillin–streptomycin–Glutamax (MeilunBio). Human embryonic kidney 293T cells were maintained at 37°C with 5% CO₂ in DMEM supplemented with 10% (v/v) FCS and 1% (v/v) penicillin–streptomycin. All cell lines used in this study were purchased from the American Type Culture Collection (ATCC).

### Mice

Male wild-type C57BL/6J mice (8–10 weeks old) were obtained from Cyagen Biosciences. Animals were group-housed (maximum five mice per cage) with ad libitum access to food and water. Housing conditions were maintained at 22°C, 50–60% relative humidity, and a 12-h light/12-h dark cycle (lights on 06:00–18:00). Mice were handled daily for 3 days prior to the onset of behavioral testing to habituate them to the experimenter. Anesthesia was administered using a safe dose of avertin. All animal procedures were performed in accordance with the Fudan University regulations on animal welfare.

### Method details

#### Vector Construction and Transfection

To construct full-length and N-terminal TAF15 overexpression cell lines, total RNA was extracted from SH-SY5Y cells using the EZ-press RNA Purification Kit and reverse-transcribed into cDNA using the 4× EZscript Reverse Transcription Mix II (with gDNA Remover). Human full-length TAF15 and the TAF15 N-term (7–99 aa) fragment were amplified by PCR (annealing temperature at 72°C). PCR products were digested with NheI and BamHI at 37°C for 3 hours and ligated into a lentiviral vector containing a CMV promoter and mCherry. For lentiviral packaging, a mixture of pMD2.G, pSPAX2, and the recombinant plasmid was incubated in DMEM for 5 minutes at room temperature, followed by the addition of 26 µL HighGene Transfection Reagent (Abclonal). The plasmid–HighGene complex was applied to 293T cells at 60–80% confluency. After 6 hours, half of the medium was replaced with fresh complete medium. Viral supernatant was collected 48 hours post-transfection by filtration through a 0.45 µm sterile filter (Millipore, HAWP04700) and used to transduce SH-SY5Y cells for 48 hours. Stably transfected cells were selected with G418 (Geneticin; 500 µg/mL) for two weeks.

To generate TAF15 knockout (KO) lines, complementary oligonucleotides encoding single-guide RNAs were annealed and digested with BsmBI at 55°C for 3 hours, then ligated into the lentiCRISPRv2 vector. Transfection procedures were identical to those described above. Following transduction, cells were selected in medium containing puromycin (1 μg/mL) for 3 days to obtain stable TAF15 KO clones. For all lentiviral overexpression experiments, lentiviral mCherry overexpression vector was used as control; for CRISPR experiments, lentiCRISPRv2 empty vector served as control.

### Immunoblotting

Cell pellets were lysed in RIPA buffer (10 mM Tris, 1 mM EDTA, 0.5 mM EGTA, 1% Triton X-100, 0.1% sodium deoxycholate, 0.1% SDS, 140 mM NaCl) supplemented with protease and phosphatase inhibitors. Protein concentrations were determined using a BCA assay kit and normalized. Primary antibodies were diluted in 1% skim milk or BSA in TBS-T (10 mM Tris-HCl, 150 mM NaCl, 0.1% Tween-20, pH 7.6) and incubated overnight at 4°C. After washing in TBS-T, HRP-conjugated secondary antibodies (anti-mouse or anti-rabbit) were applied for 1 h at room temperature. Proteins were visualized using Tanon™ Femto-sig ECL (Tanon, 180-506) and imaged with a VILBER FUSION FX Spectra system. Densitometry and quantification were performed using ImageJ.

### Nucleus-Cytoplasm Separation

Cells at 60–80% confluency were trypsinized, pelleted, and resuspended in 300 µL hypotonic buffer (20 mM Tris-HCl, 10 mM NaCl, 3 mM MgCl2, pH 7.4) containing protease inhibitors. Samples were gently triturated and incubated on ice for 20 min. 15 µL of 10% (v/v) NP-40 in PBS was added, and the samples were vortexed for 10 seconds. Centrifugation was performed at 4,000 rpm for 10 minutes at 4°C. The supernatant (cytoplasmic fraction) was collected, and the pellet was resuspended in 50 µL hypotonic buffer plus 50 µL RIPA buffer, followed by lysis on ice for 10 minutes (nuclear fraction).

### Quantitative RT–PCR

Total RNA was isolated with the EZ-press RNA Purification Kit and reverse transcribed with 4× EZscript Reverse Transcription Mix II (EZB-RT2). Quantitative PCR was performed using a SYBR Green detection system (Roche LC480). Relative expression was calculated by the 2^^−ΔΔCt^ method and normalized to the housekeeping gene GAPDH.

### Thioflavin T (ThT) Staining

Cells grown on coverslips to 60–80% confluency were fixed with 4% PFA for 20 minutes, permeabilized with 0.3% (v/v) Triton X-100 in PBS for 20 minutes, and blocked with 5% BSA in PBS for 1 hour. Coverslips were incubated with 1% (w/v) ThT in PBS for 10 minutes at room temperature, washed thoroughly with PBS-T, counterstained with DAPI and mounted. Images were acquired on a Nikon NSPARC spatial array microscope, and images were processed with NIS-Elements Viewer v5.22.

### CCK-8 Assay

Cell viability was measured with the CCK-8 kit according to the manufacturer’s protocol. Cells were seeded at 3,000 cells/well in 96-well plates with five technical replicates per group. After 72 hours, medium was replaced with 110 µL detection solution (10 µL CCK-8 reagent in 100 µL DMEM). Following 3 hours of incubation at 37°C and 5% CO₂, cell morphology was inspected under a microscope, absorbance was measured at 450 nm.

### Reactive Oxygen Species (ROS) Live Cell Flow Cytometry

Intracellular ROS levels were measured using DCFH-DA. Cells cultured in 6-well plates to 60–80% confluence were incubated with 10 μM DCFH-DA at 37°C in 5% CO₂ for 30 min protected from light. After two washes with PBS to remove excess dye, adherent cells were trypsinized to produce single-cell suspensions and analyzed on a flow cytometer (Becton, Dickinson and Company) using the FITC channel.

### snRNA-seq Analysis

Single-nucleus RNA-seq raw data and metadata from FTLD and ALS patients were obtained from Synapse (syn51105515). Data processing was performed in R (v4.5.2) using the Seurat package (v5.3.0). After integration and annotation using Seurat’s standard workflow in RStudio (v2025.09.02+418), downstream analyses were performed for specific cell types, including excitatory and inhibitory neurons.

### Bulk RNA Sequencing and Analysis

Total RNA was extracted with the EZ-press RNA Purification Kit and submitted to Novogene for sequencing. mRNA was enriched using oligo(dT) magnetic beads, fragmented, and reverse transcribed with random hexamer primers to synthesize first-strand cDNA; second-strand cDNA was synthesized using dUTP. After end repair, A-tailing, adapter ligation, fragment selection, USER digestion, amplification and purification, strand-specific libraries were generated and pooled according to effective concentration and desired output, then sequenced on an Illumina platform. Downstream enrichment analyses were performed in R using ggplot2 (v3.5.2), clusterProfiler (v4.8.3) and ComplexHeatmap (v2.16.0). Protein–protein interaction (PPI) networks were analyzed using STRING (v12.0) and visualized in Cytoscape (v3.10.3).

### Stereotaxic AAV injection

All stereotaxic injections were performed in male C57BL/6J mice (8–10 weeks old). Mice were anesthetized with a safe dose of avertin and maintained on a heating pad at 37°C during and after surgery. Stereotaxic coordinates were determined with reference to Paxinos and Franklin’s The Mouse Brain in Stereotaxic Coordinates, 5th edition (mPFC: AP +1.8, ML ±0.45, DV −2.1 relative to bregma). AAV9-EGFP and AAV9-TAF15-3×FLAG-EGFP (FUBIO, 1×10¹³ vg/mL) were injected slowly using an automated stereotaxic apparatus (RWD, 71001-S). The needle was retained for 10 minutes post-injection, the scalp was sutured and disinfected, and animals were monitored post-operatively until recovery. Mice were retained for subsequent experiments only if body weight and general health remained stable.

### Immunofluorescence

Mice were perfused with ice-cold PBS followed by 4% PFA. Brains were post-fixed overnight in 4% PFA, cryoprotected in 30% sucrose, and embedded in Tissue Tek (Sakura). Sections (20 µm thick) were blocked with 5% BSA and 0.5% Triton X-100 in PBS, then incubated with primary antibodies (1:1000) for 48 hours at 4°C. Alexa Fluor 647-conjugated secondary antibodies were applied for 2 hours at room temperature. Nuclei were counterstained with DAPI. Images were acquired on a Nikon NSPARC spatial-array microscope and processed using NIS-Elements Viewer v5.22.

### TUNEL Staining

Brain sections were permeabilized in PBS with 0.3% (v/v) Triton X-100 for 30 min at room temperature, equilibrated in Equilibration Buffer for 30 min, and positive controls were treated with DNase I (10 U/μL) for 10 min at room temperature. All samples were then incubated in TUNEL reaction mix (Equilibration Buffer, YSFlourTM640-12-dUTP Labeling Mix and Recombinant TdT) at 37°C for 1 h in the dark, washed twice with PBS-T and mounted. Images were acquired on a Nikon NSPARC spatial-array microscope and processed in NIS-Elements Viewer v5.22.

### Congo red Staining

Immerse brain sections in a 1% (w/v) Congo red solution prepared in 80% (v/v) ethanol and stain at room temperature for 15 min. Then transfer the sections to a differentiation solution of 0.1% (w/v) sodium hydroxide in 80% (v/v) ethanol for 10 s, and rinse in warm ddH₂O for 2 min. Re-stain the sections in Carazzi’s hematoxylin stain solution for 1 min, then rinse in warm ddH₂O for 2 min. Dehydrate and clear the sections by sequential immersion in 70%, 80%, 90%, 95% and 100% ethanol, followed by xylene, for 5 min each. Mount with neutral gum (neutral balsam) and image under bright-field microscopy.

### Open Field Test

Mice were placed in the lower-left corner of a 40×40 cm white arena and allowed to explore freely for 10 minutes. Behavior was recorded and analyzed using SuperMaze software (Softmaze, Shanghai). The central zone was defined as a 20 × 20 cm square; measures of total locomotion and center exploration were used to assess locomotor ability and anxiety-like behavior.

### Light-Dark Transition Test

The apparatus comprised a dark compartment (18 × 27 × 27 cm; enclosed; low illumination) and a light compartment (27 × 27 × 27 cm; open; higher illumination) connected by an opening. Animals were placed in the center of the light compartment facing away from the opening and allowed to explore for 10 min. Behavioral activity was recorded and analyzed with SuperMaze. An “entry” was scored when all four limbs entered a compartment.

### Y-Maze Spontaneous Alternation Test

The Y-maze consisted of three identical arms (35 cm long × 8 cm wide × 15 cm high) arranged at 120° angles. Mice were placed at the distal end of a designated start arm and allowed to explore freely for 10 min. Arm entries were recorded by SuperMaze. An entry was defined only as all four paws entering an arm. Spontaneous alternation was defined as consecutive entries into three different arms.

### Novel Object Recognition Test

Two identical cylindrical objects were placed at equivalent positions near the arena edges in a 40 × 40 cm box. Mice were placed facing away from the objects at equal distance and allowed to explore for 10 min (familiarization). One hour later, mice were reintroduced into the arena where one familiar object had been replaced by a novel object different in color, material and height. Interaction with novel versus familiar objects was recorded by SuperMaze to assess working memory.

### Cued Fear Conditioning Test

Each mouse was placed individually in a sound-attenuating fear-conditioning chamber with a metal grid floor; background noise at 50 dB was provided throughout. Training (day 1) began with a 2-min habituation period followed by three tone–shock pairings (tone: 3 kHz, 80 dB, 30 s; shock: 0.5 mA, 2 s delivered to the hind paw; tone and shock co-terminated). Inter-trial interval was 1 min. Animals remained in the chamber for 1 min after the final pairing before being returned to their home cage. Testing (day 2) consisted of a 3-min “cue-off” period with background noise only, followed by a 3-min “cue-on” period during which the 3 kHz tone was presented. Behavior was recorded and analyzed with ANY-Maze software. Freezing time during the two phases was compared to evaluate fear memory retrieval and regulation.

### Quantification and Statistical Analysis

Animals were randomly assigned to experimental groups using a pre-generated randomization list in Excel. All analyses were performed by investigators blinded to treatment group. Sample sizes or the number of biological replicates (n) are reported in figure legends. Two-group comparisons were performed using two-tailed unpaired Student’s t-tests. Comparisons among three or more groups were performed by one-way ANOVA or Brown-Forsythe and Welch ANOVA tests with appropriate post hoc tests. Pearson correlation was used to assess relationships between two continuous variables; R and p values are provided in figures. Corrected p-values or p < 0.05 were considered statistically significant. Statistical analyses were performed using GraphPad Prism v10.1.2. Data are presented as mean ± SEM.

**Supplemental information:** Figure S1-S3

---

**Figure S1.**
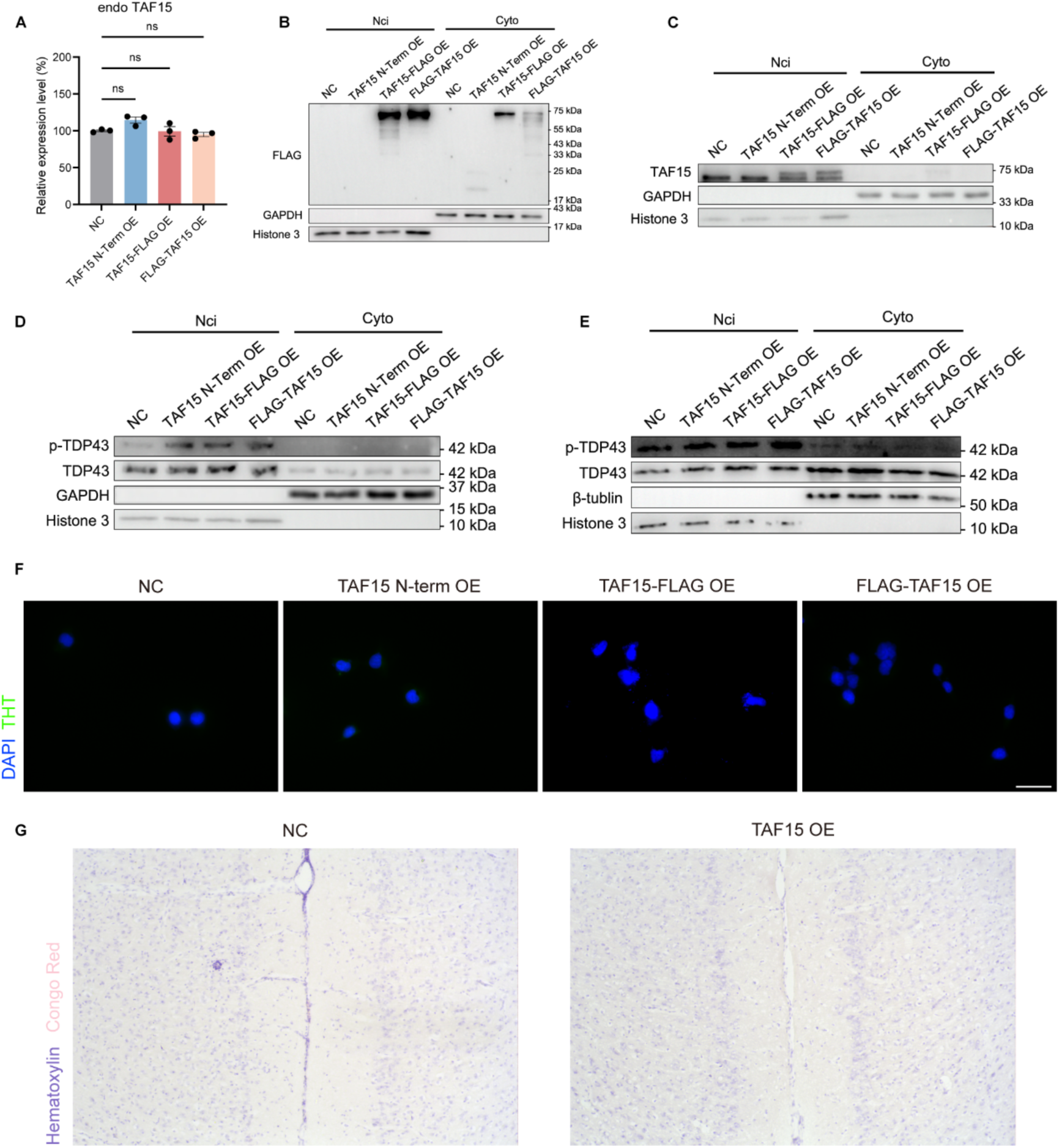
Subcellular localization analysis and assessment of amyloid formation in TAF15-overexpressing models. (A) Relative endogenous TAF15 mRNA expression in control mCherry-overexpressing cells, TAF15 N-term OE, TAF15-FLAG OE and FLAG-TAF15 OE cells (*n* = 3/group). Statistical comparisons were performed by one-way ANOVA with Dunnett’s post hoc test. (B) Representative immunoblot of nuclear-cytoplasmic fractionation (loading ratio, nuclear: cytoplasmic = 1:1) probed for FLAG, GAPDH (cytoplasmic marker), and Histone 3 in control, TAF15 N-term OE, TAF15-FLAG OE, and FLAG-TAF15 OE cells. (C) Representative immunoblot of nuclear-cytoplasmic fractionation (loading ratio, nuclear: cytoplasmic = 1:1) probed for TAF15, GAPDH, and Histone H3 in the indicated cell lines. (D) Representative immunoblot of nuclear-cytoplasmic fractionation (loading ratio, nuclear: cytoplasmic = 1:1) probed for phosphorylated TDP-43 (pTDP-43), total TDP-43, GAPDH, and Histone H3 in the indicated cell lines. (E) Representative immunoblot of nuclear-cytoplasmic fractionation (loading ratio, nuclear: cytoplasmic = 1:15) probed for pTDP-43, total TDP-43, β-actin, and Histone H3 in the indicated cell lines. (F) Assessment of amyloid formation by ThT staining in control, TAF15 N-term OE, TAF15-FLAG OE, and FLAG-TAF15 OE cells. (G) Assessment of amyloid formation by Congo red staining in mPFC of control and TAF15 OE mice.

**Figure S2.**
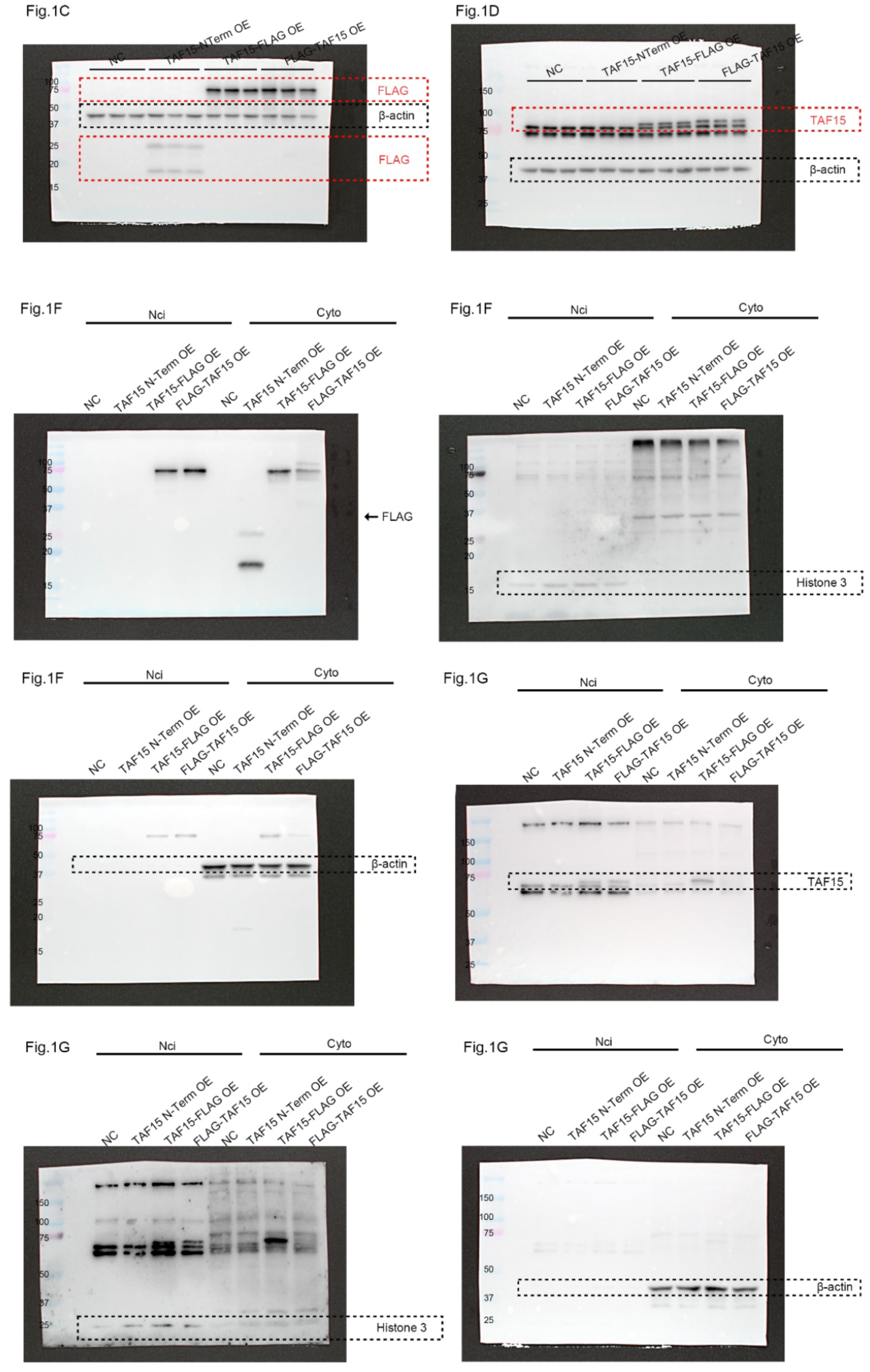
Uncropped scans of Western blots related to indicated figure.

**Figure S3.**
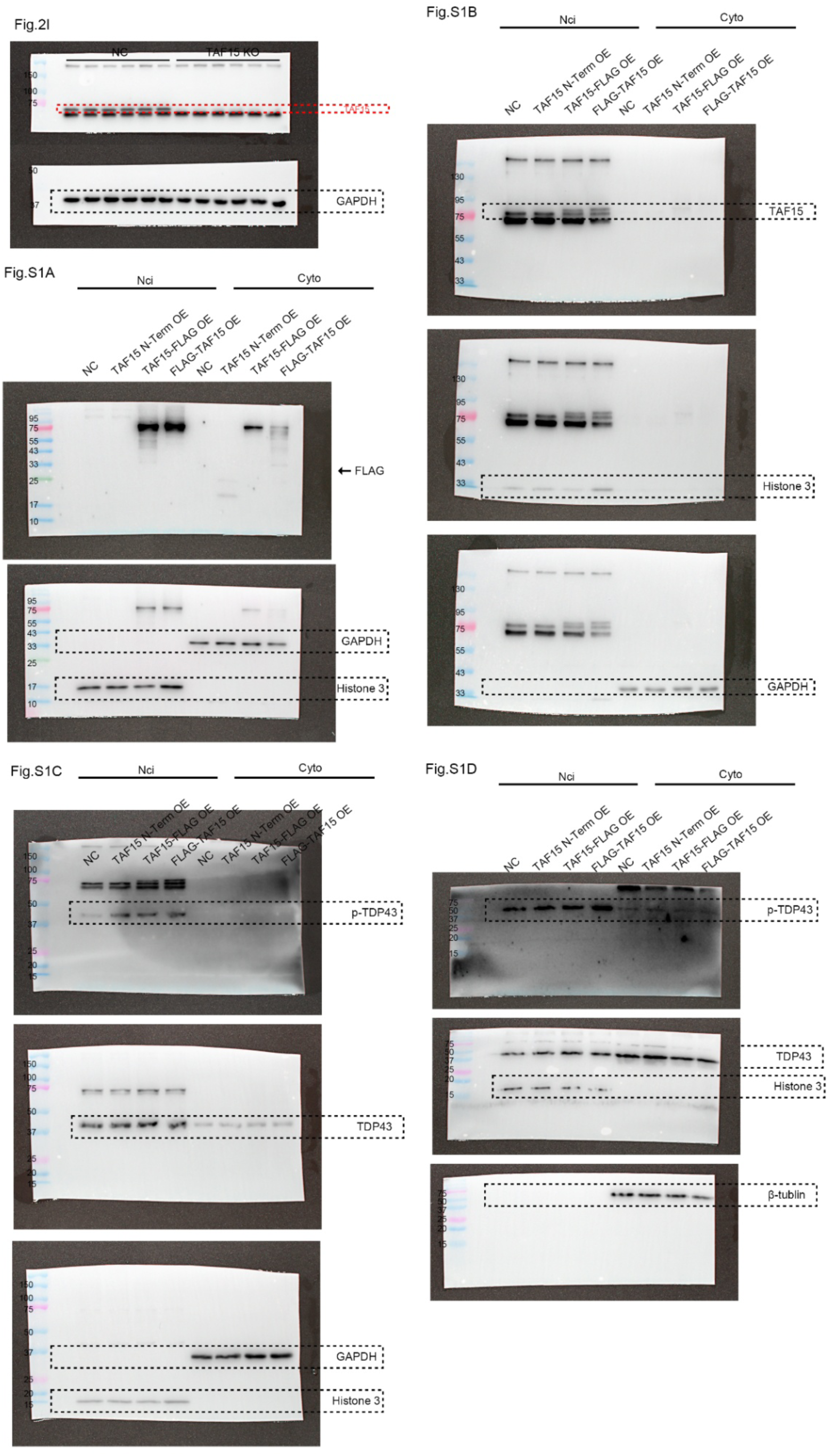
Uncropped scans of Western blots related to indicated figure.

## Notes

### Competing Interest Statement

The authors have declared no competing interest.

